# Intestinal myofibroblasts regulate intestinal epithelial cell plasticity via YAP/TAZ

**DOI:** 10.1101/2022.10.07.511327

**Authors:** Agnieszka Pastuła, Klaus-Peter Janssen, Katja Steiger, Julia Slotta-Huspenina, Richard A. Friedman, Stefanie M. Hauck, Mauricio J. A. Ruiz-Fernandez, Maximilian Fottner, Martin Borgmann, Ursula Ehmer, Roland M. Schmid, Timothy C. Wang, Michael Quante

## Abstract

Intestinal stromal cells play a key role as the crypt niche cells during epithelial homeostasis and tumor initiation. However, the underlying cellular and molecular mechanisms remain unclear. We developed various types of three-dimensional (3D) tissue culture models to culture small intestinal myofibroblasts (SI MFs) together with enteroids. SI MFs significantly enhanced self-renewal, lumen formation and survival of enteroids, that was mediated via a paracrine mechanism in a Wnt-independent manner. Such co-cultured enteroids resembled SI organoids derived from Apc+/1638N tumors. Microarray analysis showed upregulation of genes associated with YAP signaling in enteroids co-cultured with SI MFs, which was confirmed by protein quantification by mass spectrometry and could be correlated with findings from human colorectal tumor specimens. Mass spectrometric analysis of conditioned media and inhibitor studies pointed to a role for TGF-β in the SI MF-SI epithelium cross-talk. Altogether, utilizing different 3D stroma-epithelium co-culture models, we demonstrate here that SI MFs have the potential to induce a tumor-like phenotype in the intestinal crypts via a paracrine mechanism, that involves YAP and TGF-β, but not canonical Wnt signaling.

## INTRODUCTION

Epithelial cells are present in the majority of our organs. Depending on the organ, epithelial cells can play a barrier function, absorb nutrients, or perform secretory functions. Epithelial cells regenerate regularly under homeostatic conditions during physiological cell turnover in an adult organism and can be a source of a tumor. Epithelial cells exhibit plasticity that is crucial during embryogenesis, homeostatic conditions, and injury. Additionally, epithelial cell plasticity is implicated in diseases such as cancer and therapeutic escape. Nevertheless, the mechanisms by which epithelial cell plasticity is regulated are not fully understood.

The tissue microenvironment is defined as the tissue surrounding that is composed of neighboring cells, extracellular matrix, and soluble factors. The tissue microenvironment regulates many cellular processes such as immune response, cellular migration, metabolism, proliferation, and differentiation (Christo et al., 2021; Komohara and Takeya, 2017; Belli et al., 2018). The stem cell niche appears as a highly specialized tissue microenvironment in an adult organism and is believed to regulate stem cell self-renewal and protect stem cells against terminal differentiation (Voog and Jones, 2010). One of the tissues with the highest cell renewal rates during homeostatic conditions is the intestinal epithelium. Its constant state of regeneration that is fueled by the intestinal stem cells (ISCs) (Barker et al., 2012, 2007; Tan et al., 2021)., renders it prone to tumor formation.

Although there has been progress in understanding ISC biology (Böttcher et al., 2021; Beumer and Clevers, 2016; Zeve et al., 2022), there is still very little known about the regulatory mechanisms in the intestinal stem cell niche and its impact on tumor initiation. Paneth cells, an intestinal epithelial cell type with immune defense functions, express niche signals such as Wnt3, EGF, TGFα and the Notch ligand Dll4 (Sato et al., 2011). Nevertheless depletion of Paneth cells in mice does not significantly affect the ISCs (Durand et al., 2012), which suggests the potential role of other cell populations from the microenvironment such as stromal cells. Single cell RNA analyses of the intestine suggest existence of multiple mesenchymal cell subpopulations (Smillie et al., 2019; Fawkner-Corbett et al., 2021; Bahar Halpern et al., 2020). However, those are transcriptional cell subpopulations and for most of them functional studies are needed to decipher the precise role during homeostasis and disease. α-SMA+ myofibroblasts (MFs) are spindle*-*shaped cells that are adjacent to intestinal epithelial cells in both the villi and the crypts. Interestingly, α-SMA+ MFs are unabundant in healthy intestine, but their number is greatly increased in the tumor microenvironment (D’Arcangelo et al., 2020) suggesting that these cells expand during cancer formation. α-SMA+ MFs have been proposed as a potential therapeutic target for cancer, however the mechanisms regulating interactions of adult intestinal α-SMA+ MFs with normal intestinal epithelium remain unknown.

Research using *in vivo* and *in vitro* models showed that the stromal niche in the intestine consists of various mesenchymal cell populations: Foxl1+ mesenchymal cells and CD34+ mesenchymal cells (Aoki et al., 2016; Shoshkes-Carmel et al., 2018; Stzepourginski et al., 2017) have been demonstrated to be an important component of the intestinal stem cell niche, however the role of adult subepithelial α-SMA+ MFs (also known as pericryptal MFs)(Worthley et al., 2010) that surround the intestinal crypt, remains largely unknown.

Here, we aim to revisit the concept of α-SMA+ MFs to epithelial cell interactions in a 3D culture resembling the intestinal stem cell niche and analyze their function during intestinal tumor initiation, as heterotypic intercellular communication is distorted in tumor and the number of α-SMA+ MFs is increased during intestinal tumorigenesis (Adegboyega et al., 2002).

## RESULTS

### Intestinal MFs induce enterospheres and promote self-renewal of intestinal crypts *in vitro*

To investigate stromal-epithelial cell-cell interactions of the small intestinal (SI) stem cell niche, we utilized a 3D co-culture system combining intestinal crypts and intestinal MFs (Pastuła et al., 2016; Pastuła and Marcinkiewicz, 2018). Mouse SI crypts and SI MFs were isolated from WT C57BL/6 mice and cultured separately (Figure 1A). Cultured epithelial and stromal cells exhibited high purity as indicated by RT-PCR, with specific expression of E-cadherin in the crypts and α-SMA in the intestinal MFs (Figure 1 A). After expansion, SI MFs were combined with SI crypts and cultured in a Matrigel drop (Figure 1 A). The co-culture resulted in fast and impressive phenotypic changes regarding the growth of the SI crypts with formation of round structures with a large lumen and thin epithelial cell monolayer; such structures were defined as enterospheres (Figure 1 B). Under co-culture conditions 54% of such enterospheres were observed in contrast to 3.2% under monoculture conditions (Figure 1 C). Routine SI crypt culture requires addition of external niche factors such as R-Spondin, EGF and Noggin (Sato et al., 2009; Pastuła and Quante, 2016), which were not necessary in co-culture conditions (Figure 1 B). Without growth factors (GFs), we observed 2.3% enterospheres in the monoculture, while addition of SI MFs resulted in 53.1% of enterospheres (Figure 1 C). Interestingly, additional to SI MFs, we observed induction of enterospheres also through colon MFs, mouse gastric carcinoma associated fibroblasts (CAFs) and human cardia MFs (Figure S1). Clonogenicity assays (Figure 1 D) revealed that the presence of SI MFs increased formation of single cell-derived organoids by 4.5 ± 1.2-fold (Figure 1 E). In addition, SI MFs increased the diameter of single cell derived organoids from 48.6 ± 3.4 (monoculture) μm to 75.3 ± 3.6 μm (co-culture) (Figure 1 F). Thus, MFs influenced growth and structure of the intestinal organoids independently of external niche factors in an *in vitro* intestinal stem cell niche.

**Figure 1.**
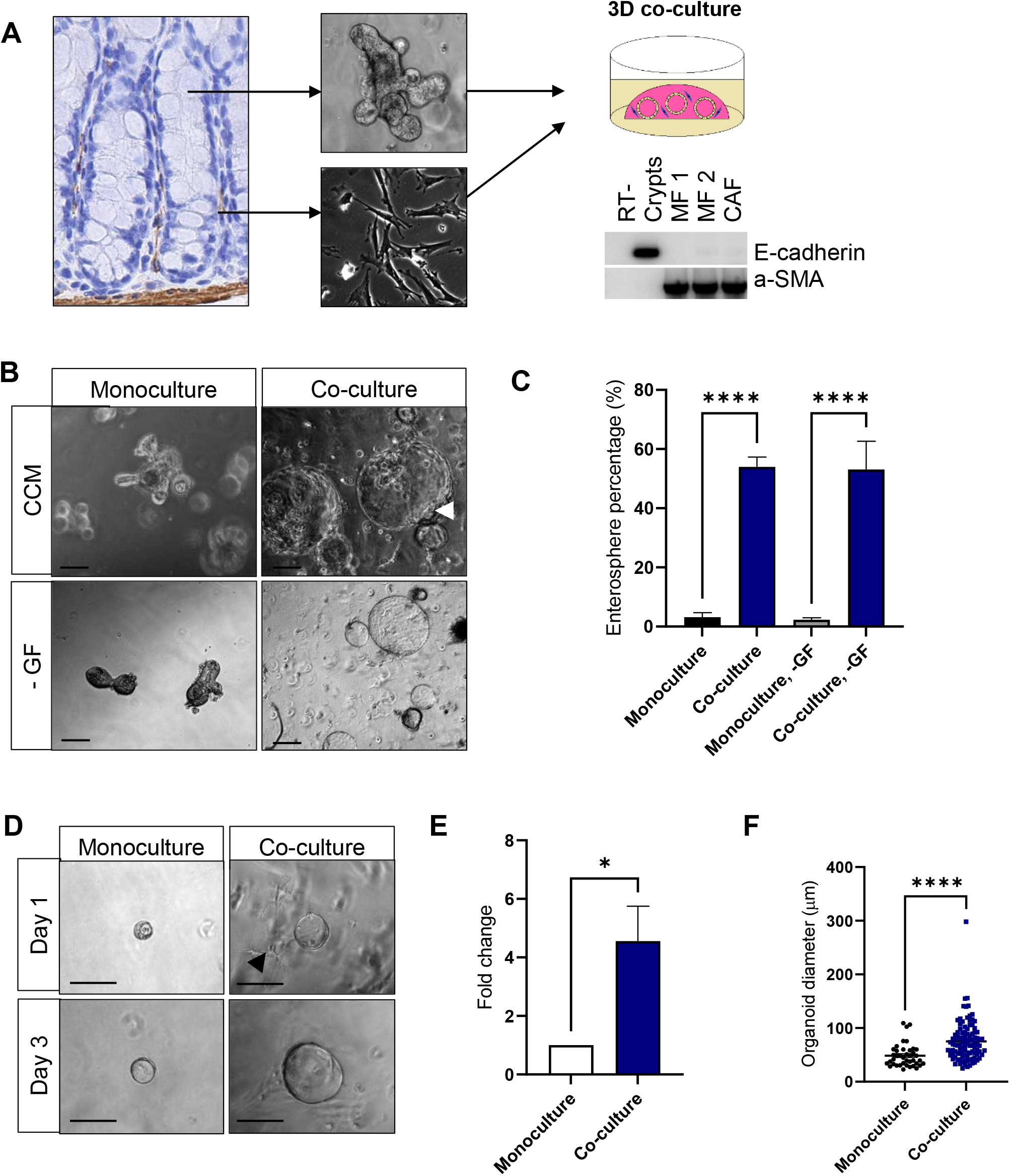
Myofibroblasts (MFs) promote self-renewal and induce enterospheres in the intestinal crypt *in vitro* independently of niche factors: R-Spondin, EGF and Noggin.**A**. Research model: small intestinal (SI) MFs and SI crypts were isolated, expanded and then combined in a 3D culture system. RT-PCR shows the purity of cultured cells. **B**. Morphology of SI organoids in the presence of SI MFs. An enterosphere is marked with an arrowhead. Scale bar, 100 μm. **C**. Quantification of enterospheres in SI organoids in culture conditions with and without growth factors (GF): R-Spondin, EGF and Noggin; mean ± SEM; one-way ANOVA, Bonferroni’
ss multiple comparison test, p < 0.0001. **D**. Morphology of single-cell derived SI organoid cultures in the presence of SI MFs. An arrowhead indicates myofibroblasts. Scale bar 100 μm. **E**. Quantification of single-cell derived SI organoids in the presence of SI MFs (day 3); mean ± SEM of three independent experiments; two-tailed t-test, p = 0.0409. **F**. Organoid diameter in single-cell derived SI organoid cultures in the presence of SI MFs (day 3); mean ± SEM, two-tailed t-test, p < 0.0001.

### Intestinal MF - intestinal epithelium interactions are specifically mediated by the mesenchymal cell-derived soluble factors

Cell-cell interactions in the niche can be either contact dependent or paracrine. It has been proposed that direct contact is needed for Wnt3-dependent interaction between Paneth cells and ISC (Farin et al., 2016). Since culturing SI crypts and SI MFs together in a Matrigel drop (Figure 1 A and B) may enable direct stromal cell-epithelial cell contact, we utilized an indirect co-culture system with a membrane (transwell) placed between MFs (top) and crypts (bottom) (Figure 2 A) to investigate whether direct stroma-epithelium contact was necessary for the formation of enterospheres. SI organoids in such a co-culture system formed enterospheres (Figure 2 B), similarly to the direct co-culture (Figure 1 B). This finding was confirmed by conditioned media experiments in which we transferred co-culture exposed media to the single crypt cultures, which revealed that the MF-conditioned medium (MCM) alone was sufficient to initiate formation of enterospheres in the SI crypts *in vitro* (Figure 2 B). In addition to the induction of enterospheres, MFs significantly increased the diameter of organoids. While in the control organoids (monoculture/untreated) average organoid diameter was 75.4 μm, in the organoids from the indirect co-culture it was 162.9 μm, and in the organoids treated with MCM it was 147 μm (Figure 2 C), suggesting that SI MFs regulate growth of crypt cells by a paracrine mechanism. Crypt cultures treated with MCM there were 34.4% enterospheres, in contrast to control crypt cultures that contained only 1.3 enterospheres (Figure 2 D). Thus, the stromal cell secretome specifically induces enterospheres in WT SI crypts. To investigate this phenomenon further, we treated SI organoids with MCM for 4h and found that MCM caused a 3.8-fold increase of the lumen area in SI organoids (Figure 2 E and F). Taken together, MFs induced morphological alterations in intestinal crypts *in vitro* by induction of enterospheres and promoted lumen formation via soluble factors in the absence of external niche factors.

**Figure 2.**
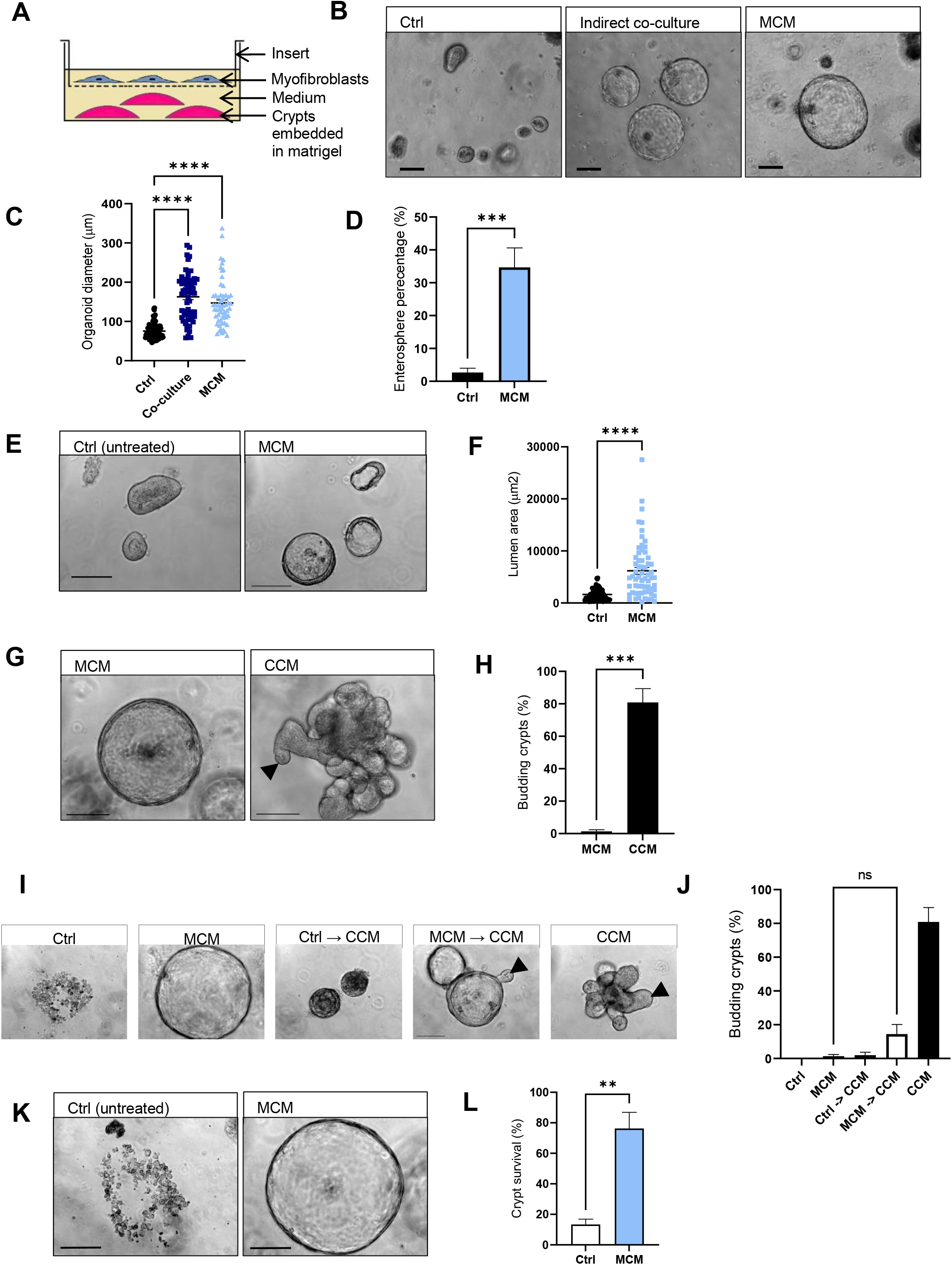
Heterotypic stroma-epithelium interactions in the intestinal stem cell niche are mediated by stromal cell-derived soluble factors.**A**. Scheme of the indirect stroma-epithelium co-culture system. **B**. Morphology of small intestinal (SI) organoids either co-cultured with SI myofibroblasts (MFs) or incubated with SI MF conditioned medium (MCM). Scale bar, 100 μm. **C**. Organoid diameter in SI organoids from indirect co-cultures and cultures treated with MCM. One-way ANOVA, Bonferroni’
ss multiple comparison test, p < 0.0001. **D**. Quantification of enterospheres in crypt cultures treated with MCM, mean ± SEM of three independent experiments: t-test, p = 0.0002. **E**. 4 h-exposure to the MF-derived soluble factors: MCM induces lumen formation in the intestinal crypts *in vitro* in the absence of external niche factors R-Spondin, EGF and Noggin. Scale bar, 100 μm. **F**. Analysis of lumen area in the SI organoid cultures treated with MCM for 4 h. Two-tailed t-test, p < 0.0001. **G**. Stromal cell derived soluble factors (MCM) do not promote budding of intestinal crypts *in vitro*. An arrowhead marks a bud in the organoid. Scale bar, 100 μm. **H**. Quantification of budding organoids, two-tailed t-test, p = 0.0007. **I**. Myofibroblast-induced epithelial spheres start generating buds upon addition of culture medium containing EGF, Noggin and R-Spondin (crypt complete medium, CCM). Scale bar, 100 um. An arrowhead indicates buds. **J**. Quantification of budding crypts upon treatment with MCM and medium switch to CCM. Three independent experiments. One-way ANOVA. Ns, not significant. **K**. Stromal-cell derived soluble factors (MCM) promote intestinal crypt survival *in vitro* in the absence of external niche factors: R-Spondin, EGF and Noggin. Scale bar, 100 μm. **L**. Analysis of organoid survival, two-tailed t-test, p = 0.0049. For all experiments in B – L, the MCM was generated from the cultures of SI MF only.

Next, we asked whether MF-derived soluble factors had any impact on a SI organoid differentiation phenotype. Previously described mini-gut cultures that were maintained in the presence of external niche factors (R-Spondin, EGF and Noggin) but without the stromal niche, were characterized by the presence of numerous buds as a surrogate for differentiation (Sato et al., 2009; Pastuła and Quante, 2016). We found that SI organoids incubated with MCM (Figure 2 G) had only 1.4 ± 1 % budding crypts, in contrast to SI organoids cultured in the presence of EGF, Noggin and R-Spondin (Figure 2 G, CCM) that contained 80.9 ± 8.5 % budding crypts (Figure 2 H), suggesting that SI MF-derived soluble factors induced appearance of enterospheres and promoted lumen formation but decreased crypt budding.

To investigate whether MF-induced enterosphere phenotype is reversible, the crypts were firstly treated with MCM; and then the medium was switched to crypt complete medium (CCM), that contains EGF, Noggin and R-Spondin. Interestingly, we observed that CCM can induce formation of buds in MF-induced enterospheres (Figure 2 I). Quantification of budding crypts revealed that after addition of CCM to MF-induced enterospheres 14.4 % of crypts formed buds (Figure 2 J, MCM → CCM), in contrast to control enterosphere cultures, in which 1.4 % of budding crypts were identified (Figure 2 J, MCM).

One of the known functions of the stem cell niche is to provide signals regulating survival of stem cells (Jones and Wagers, 2008). To investigate whether SI MF-derived soluble factors could promote survival of crypt cells, we cultured SI organoids for 3 days in the absence of EGF, R-Spondin and Noggin. Organoids lost the capability to form any epithelial layer (such structures were defined as nonviable organoids) (Figure 2 K); only 13.4 ± 3.5% of SI organoids remained viable, in contrast to organoid cultures with MCM 76.3 ± 10.6 % (Figure 2 L), suggesting that SI MF secretome also provides survival signals for the intestinal epithelium.

### MFs induce poorly differentiated and proliferative phenotype in SI organoids additional to cell intrinsic Wnt signaling

To characterize the phenotype of SI crypts cultured in the presence of MF-derived niche factors we found that expression of Lgr5, an intestinal stem cell marker, was not affected (Figure S2 A). However, Sox-9 (marker of progenitor cells in the intestine) and CD44 (cancer stem cell marker) expression was increased 3.6-fold (p = 0.0348) and 4.3-fold (p = 0.0069), respectively (Figure S2 A). In addition, we observed elevated expression of proliferation markers, such as cyclin D1, but not differentiation markers mucin-2 and cryptidin-5 (Figure S2 A). Together with the clonogenicity assay (Figure 1 D - F), these data suggest that MF-derived niche factors promote cellular proliferation, and rather de-differentiation and not differentiation in the intestinal cell lineage. To confirm the differentiation status, we quantified the number PAS+ cells (Goblet cells) in SI organoids co-cultured with SI MFs and observed that in the presence of MFs the number of PAS+ cells was reduced (Figure S2 B).

Similar niche signals regulate normal adult stem cells and tumor-initiating cells (Giancotti, 2013). MFs provide a link between the normal adult stem cell niche and the tumor niche in the intestine (Pastuła and Marcinkiewicz, 2018). As we observed that MF-conditioned medium upregulates CD44 (a cancer stem cell marker) in the crypts (Supplementary figure S2 A); this prompted us to test whether MFs have the potential to regulate cellular processes, such as proliferation and differentiation, that are deregulated during tumor initiation. We compared SI organoids from co-culture (or organoids treated with MCM) with adenoma (Apc^+/1638N^) SI organoids. Phase contrast images, Ki-67 staining, and PAS staining revealed that both the co-culture/ MCM-treated enteroids and adenoma enteroids were enriched for enterospheres, contained proliferating cells in the absence of EGF, R-Spondin and Noggin; and exhibited decreased number of goblet cells, as compared to the intestinal organoid monoculture (Figure 3 A). To investigate the impact of intestinal MFs on the gene expression in the normal intestinal epithelium and whether MFs could induce the expression of genes associated with tumor initiation, we performed microarray analysis of the crypts from co-cultures with SI organoids in the presence of SI MFs and in the absence of external niche factors (Figure 3 B) as well as of adenoma organoids cultured without the external niche factors. Microarray analysis revealed that in the co-culture 169 genes were significantly upregulated and 36 genes downregulated when compared to the monoculture. Venn diagram of differentially expressed genes showed that 123 (75%) upregulated genes and 31 (86%) downregulated genes in the co-culture overlapped with the equivalent genes in the adenoma organoids (tumor-initiation gene signature; Figure 3 C and S2 C). *In silico* gene expression analysis using DAVID bioinformatics revealed the involvement of genes associated with important cancer associated signaling pathways, such as pathways associated with cell cycle, focal adhesion, ECM-receptor pathway (Figure 3 C). Interestingly, we discovered that MFs can regulate expression of genes involved in cellular metabolism (Figure 3 C), and altered cellular metabolism was previously found to be linked to intestinal tumor initiation (Bensard et al., 2019).

**Figure 3.**
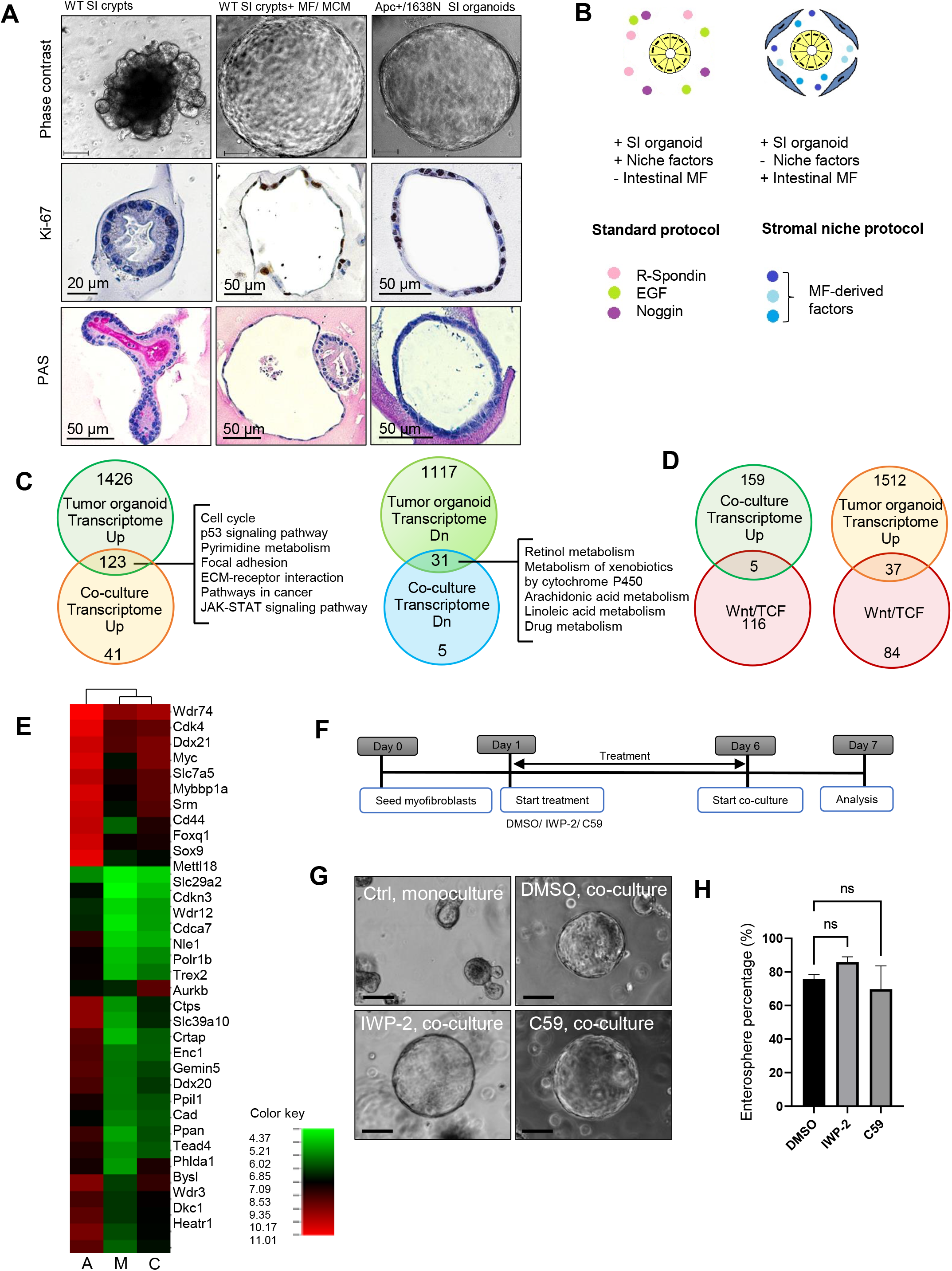
Small intestinal (SI) myofibroblasts (MFs) induce poorly differentiated phenotype in the SI organoids by other mechanism than the canonical Wnt.**A**. Morphology (scale bar 100 μm), Ki-67 staining and PAS staining of wild type SI organoid monoculture, wild type SI organoids co-cultured with SI myofibroblasts or MCM, and tumor Apc^+/1638N^ organoids. **B**. Schematic diagram of the short-term small intestinal (SI) organoid culture in the presence of stromal niche for the microarray analysis. Previously published protocol involved SI organoid monoculture in the presence of niche factors: R-Spondin, EGF and Noggin. To study the stromal niche we optimized a protocol that involves SI organoid culture in the presence of SI MFs and in the absence of niche factors R-Spondin, EGF and Noggin. **C**. Venn diagram and pathway analysis of the overlapping genes (’
s’tumor-initiation signature’’) between tumor Apc^+/1638N^ organoids and wild type SI organoids co-cultured with SI myofibroblasts. Up, upregulated. Dn, downregulated. **D**. Common genes for the Wnt/TCF signature and genes upregulated in the co-culture/ tumor Apc^+/1638N^ enteroids. **E**. Heatmap for the genes from the intestinal Wnt/TCF signature. A, tumor Apc^+/1638N^ enteroids. M, wild type organoid monoculture. C, wild type organoid co-culture. **F**. Experimental outline for the Wnt inhibition experiment. **G**. Morphology of wild type enteroids in the co-culture with myofibroblasts and upon blockade of Wnt secretion by small molecule inhibitors IWP-2 and C59. Scale bar 100 μm. **H**. Enterosphere percentage in the co-cultures after blockade of Wnt secretion. Ns, not significant.

Next, we asked which molecular mechanism was involved in the MF-epithelial crypt cross-talk. Of note, in the crypts from the co-culture there were only 5 overlapping genes (4.1%) (Figure 3 D), in contrast to 37 genes (30.6 %) of the list of modified genes in adenoma organoids that were overlapping with the intestinal Wnt/TCF signature, which is thought to be crucial for the maintenance of ISC (Van der Flier et al., 2007) (Figure 3 D and S2 C). Of note, the intestinal Wnt/TCF signature in the crypts from the co-culture clustered more with the monoculture than with the adenoma organoids (Figure 3 E), suggesting that the MF-crypt cross-talk in our model could be mediated by other mechanism than Wnt signaling.Still, RT-PCR showed that SI MFs expressed *Wnt5a, Wnt9a* and other niche factors such as *Bmp4, Grem1, Grem2, Chrdl1, Fstl3* and *FGF2* (Figure S2 D). For Wnt inhibition, SI MFs were treated either with IWP-2 or C59 to inactivate porcupine (Chen et al., 2009; Proffitt et al., 2013), and then cultured with SI organoids (Figure 3 F). Blockade of Wnt secretion altered neither the morphology of enterospheres (Figure 3 G) nor enterosphere percentage (Figure 3 H). In addition, expression of *Axin2*, as a feedback loop mechanism upon activation of Wnt signaling (Yan et al., 2001; Jho et al., 2002; Lustig et al., 2002), was not increased upon treatment of SI organoids with MCM (Figure S2 E). These data suggest that stroma-epithelium interactions in the reconstructed intestinal stem cell niche are likely mediated by a mechanism other than canonical Wnt signaling.

### TGF-β and YAP regulate the MF-induced, proliferative enterosphere phenotype in the intestinal stem cell niche

Proteomic profiling of the supernatant from mouse and human MF-SI organoid co-culture by mass spectrometry (Figure 4 A) identified 51 mouse proteins (Figure S3 A) and 22 human proteins (Figure S3 B) that were significantly upregulated when compared to SI organoid monoculture; these were mainly enriched for extracellular matrix proteins (Figure S3 C and D). Especially proteins associated with TGF-β signaling (and TGF-β interacting partners) such as TGFBI, THBS, CTGF, Lrg1 and decorin (Figure S3 A, B), suggested the involvement of the TGF-β pathway in the stromal-epithelium cross-talk. SI MFs expressed *Tgfb1, Tgfb2* and *Tgfb3*, whereas SI crypts cultured *in vitro* expressed *Tgfbr1, Tgfbr2* and *Tgfbr3* (Figure 4 B). To investigate whether TGF-β was involved in the stromal-epithelium cross-talk we inhibited TGF-β signaling utilizing the TGF-β receptor-1 and −2 dual inhibitor, LY2109761 (Melisi et al., 2008). Indeed, SI MF-SI organoid co-culture incubated with LY2109761 had reduced diameter (145.5 ± 5 μm) when compared to the DMSO-treated co-culture (236.5 ± 12 μm) (Figure 4 C and D). In addition, upon LY2109761 treatment enterosphere percentage was decreased from 43.2 % (DMSO-treated co-culture) to 7.3 % (LY2109761-treated co-culture) (Figure 4 E), which suggests that TGF-β signaling is critical for the enterosphere formation.

**Figure 4.**
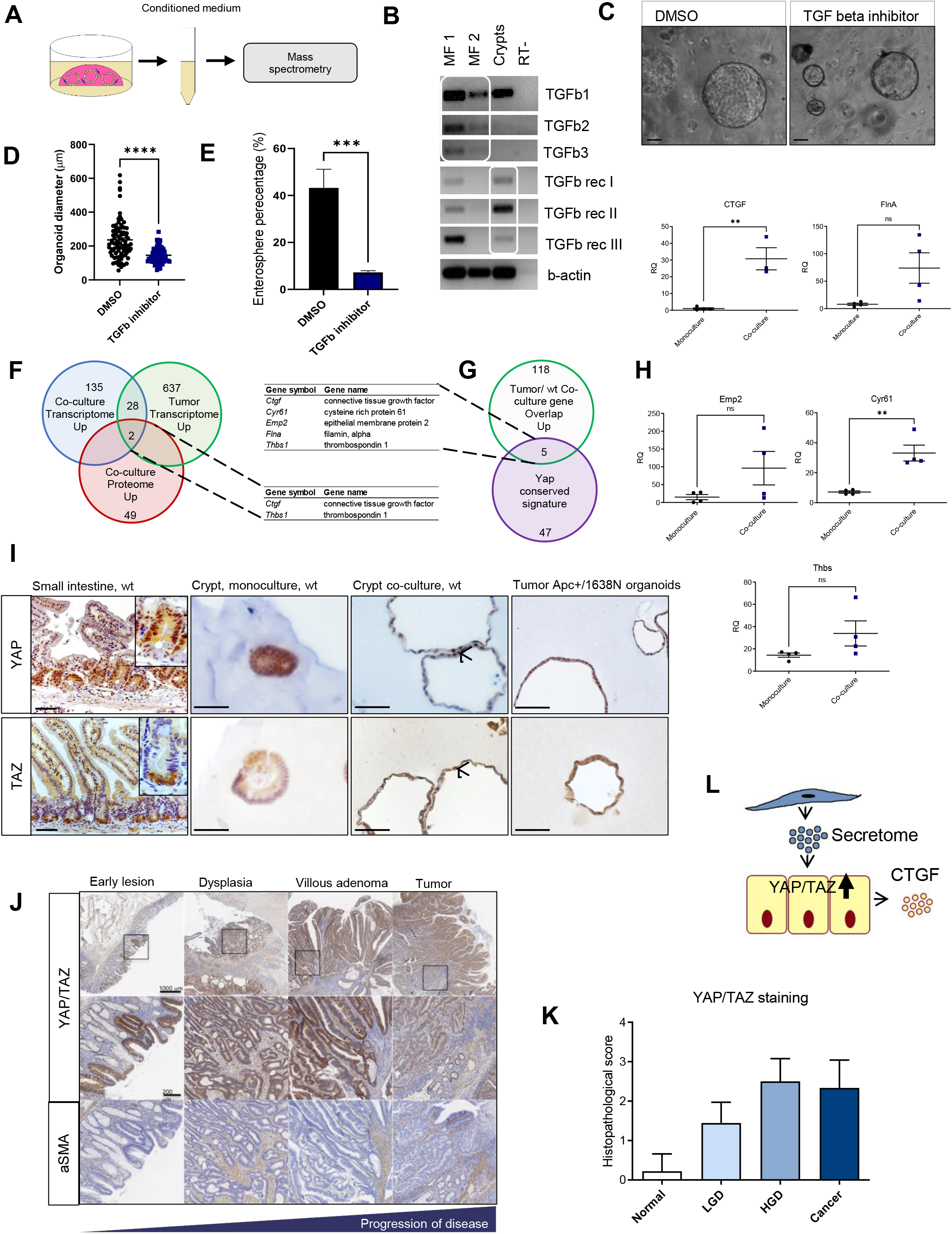
YAP/TAZ provides a link between the normal stem cell niche and the tumor initiation niche.**A**. Experimental setup for the analysis of secreted protein(s) that affect local stem cell niche and influence the cross-talking between epithelial cell and mesenchymal cell. Enteroids were co-cultured either with murine SI myofibroblasts (MFs) or human cardia MFs. The supernatant was harvested for mass spectrometry analysis. **B**. RT-PCR analysis of the components of TGF-β pathway in the SI crypts and SI MFs cultured *in vitro*. RT-, negative control for reverse transcription. MF P3 and MF P23 indicate intestinal myofibroblast cultures with different passage numbers. **C**. Morphology of intestinal organoids in the co-culture with MFs upon inhibition of TGF-β by LY2109761. Scale bar 100 μm. **D**. Organoid diameter in cultures treated with LY2109761. Two-tailed t-test, p < 0.0001. **E**. Enterosphere percentage in cultures treated with LY2109761. Two-tailed t-test, p = 0.0005. **F**. Upregulated genes in the SI organoid co-culture and Apc^+/1638N^ organoids from microarray data in comparison to upregulated proteins from the co-culture. **G**. Upregulated genes common for the WT enteroids and Apc^+/1638N^ enteroids versus Yap conserved signature. **H**. Validation of microarray data: expression of CTGF (two-tailed t-test, p = 0.0031), Cyr61 (two-tailed t-test, p = 0.0028), Emp2, FlnA and Thbs in the SI organoids from the indirect co-culture, evaluated by real-time PCR. **I**. Immunohistochemical staining of YAP/TAZ of murine small intestine (SI), SI organoid monoculture, SI organoid co-culture and Apc^+/1638N^ organoids. Arrowheads indicate nuclear YAP and nuclear TAZ in SI organoid co-culture. Scale bar 50 μm. **J**. Representative pictures and **K**. Quantification of immunohistochemical staining of YAP/TAZ and α-SMA in different lesions during colorectal carcinogenesis: normal colon mucosa (n=9), adenoma with low grade dysplasia (LGD) (n=9) or high grade dysplasia (HGD) (n=4) and colon cancer (n=9), ANOVA p<0.0001. **L**. Schematic diagram of the interaction of intestinal MFs with the intestinal epithelium.

Of note, a combined analysis of the transcriptome and proteome overlapping signaling molecules revealed an upregulation of CTGF (Figure 4 F), a direct TGF-β partner (Abreu et al., 2002) and downstream target of YAP/TAZ (Lai et al., 2011), two transcriptional co-activators and critical regulators of organ size and stem cell self-renewal (Zhao et al., 2011). Interestingly, the gene overlap between adenoma organoids and WT organoids from the co-culture (Figure S2 C) contained genes from a YAP conserved signature (Cordenonsi et al., 2011) including *Cyr61, Ctgf, Emp2, Thbs* and *Flna* (Figure 4 G). Validation of microarray data confirmed that the expression of genes associated with YAP activation: *Cyr61, Ctgf, Emp2, Thbs* and *Flna*, was increased in the co-culture when compared to the monoculture (Figure 4 H), suggesting that MFs can regulate YAP/TAZ signaling. YAP/TAZ staining demonstrated that in the murine SI *in vivo* YAP/TAZ was specifically expressed in the crypt compartment, but not in the villus, representing the differentiated compartment (Figure 4 I). Consistently, we detected active (nuclear) YAP/TAZ in the crypts *in vitro*, in both WT crypt co-culture and adenoma organoids (Figure 4 I). To examine the involvement of YAP in MF-epithelium interactions in the *in vitro* intestinal stem cell niche further, we treated SI organoids simultaneously with MCM and cytochalasin D (Figure S3 E), an actin destabilizing agent, which was shown to inhibit YAP nuclear localization (Johnson and Halder, 2014): cytochalasin D-treated cultures had an average enterosphere diameter of 77.3 ± 2 μm, compared to DMSO-treated cultures with 129.8 ± 4 μm (Figure S3 F). Moreover, DMSO-treated cultures contained 65.1 % of enterospheres, while cytochalasin D-treated cultures only 24 % (Figure S3 G).

Finally, we observed a significant increase of YAP/TAZ signaling in correlation with α-SMA expressing stromal cells during colon carcinogenesis in human biopsy material representing normal colon mucosa in comparison to adenoma with low grade dysplasia (LGD, early lesion) or high-grade dysplasia (HGD) and colon cancer (n=9) (Figure 4 J, K). Altogether, our data suggest that intestinal MFs modulate intestinal epithelial homeostasis and tumorigenesis by providing secreted niche factors that can regulate YAP/TAZ in the adjacent epithelium (Figure 4 L).

## DISCUSSION

Heterotypic cell-cell interactions play an essential role during morphogenesis and epithelial tissue maintenance in an adult organism and are severely altered during many chronic diseases such as cancer. Here, we describe a role of α-SMA+ mesenchymal cells, that surround the intestinal crypt, in intercellular interactions between stromal and epithelial cells in the intestinal stem cell niche. Our data point to the YAP-TGF-β axis as a potential mechanism that links the normal stem cell niche and epithelial tumor initiation, and that is regulated by the stromal microenvironment.

We demonstrate that the stromal microenvironment regulates self-renewal and cellular differentiation in the intestinal epithelium in a 3D cell culture model mimicking the intestinal stem cell niche. α-SMA+ MF-derived factors modulate proliferation, differentiation and promote survival in the intestinal crypt culture, therefore α-SMA+ MFs can be considered as a constituent of the intestinal stem cell niche *in vitro*. Previously, Paneth cells were shown to provide crucial niche factors for ISCs (Sato et al., 2011), however deletion of Paneth cells in mice did not alter ISCs (Durand et al., 2012). Therefore, α-SMA+ MFs emerge as key regulators of ISC maintenance and regulate cellular proliferation and differentiation in the intestinal crypt. In addition, we show that interactions of α-SMA+ MF with the intestinal epithelium do not require direct cell-cell contact, in contrast to Wnt3-mediated Paneth cell – ISC interactions (Farin et al., 2016). As various populations of stromal cells such as CD34+ mesenchymal cells (Stzepourginski et al., 2017) and Foxl1+ mesenchymal cells (Aoki et al., 2016) and enteric glial cells (Baghdadi et al., 2022) have been shown to provide a niche for ISCs we would argue that Paneth cells together with different non-epithelial cell types might form a supportive niche for ISC.

The stromal microenvironment proves to induce poorly differentiated and proliferative phenotype identified in the WT SI crypts upon exposure to α-SMA+ MFs (or α-SMA+ MF-derived factors); such a phenotype resembles adenoma crypts that were derived from Apc^+/1638N^ tumors. Therefore, our study suggests that α-SMA+ MFs might be responsible for ISC maintenance and additionally capable of promoting tumor initiation in the intestinal epithelium. Importantly, α-SMA+ MF are one of the most abundant non-malignant cell types in the tumor microenvironment and a lot of malignant tumors are characterized by a large stromal contribution. Moreover, α-SMA+ MFs can be potential prognostic factors in colorectal cancer (Tsujino et al., 2007) and are found early on during gastrointestinal tumorigenesis (Quante et al., 2011). Although, there is a large body of experimental evidence that MFs promote tumor growth, angiogenesis and tumor cell invasion (Orimo et al., 2005; De Wever et al., 2008; Quante et al., 2011), the role of α-SMA+ MFs during tumor initiation remains largely unknown. Since α-SMA+ MFs can induce phenotypic convergence of crypts from the co-culture to tumor crypts, we argue that intestinal α-SMA+ MFs have the potential to contribute to epithelial tumor initiation. Interestingly, the crypt-MF interactions resembling a potential tumor initiation process in our model could be mediated via a paracrine mechanism that is independent of canonical Wnt signaling. This explanation is consistent with the study of Roman *et al*. (San Roman et al., 2014), in which deletion of porcupine in subepithelial MFs did not significantly affect intestinal crypt homeostasis. Furthermore, it could be that in our MF-intestinal epithelium co-culture system Wnt signaling is transiently inhibited by Yap, as observed by Gregorieff et al. (Gregorieff et al., 2015).

In this study, we identified TGF-β and YAP signaling to be involved in a potential molecular mechanism of the stroma-epithelium cross-talk in the intestinal stem cell niche. Our mechanistic studies of α-SMA+ MF – epithelial cross-talk point to the complexity of a stromal secretome that includes different ECM proteins and TGF-βI. TGF-β signaling plays and important role during cancer progression, and was shown to be an important component of the stem cell niche (Oshimori and Fuchs, 2012). Of note, in the intestinal stem cell niche TGF-β gradient is observed along the crypt-villus axis (Pelton et al., 1991).

Gene expression analysis demonstrate that α-SMA+ MFs stimulate the expression of YAP transcriptional targets in our 3D *in vitro* model. YAP is associated with stem cell traits (Lian et al., 2010) and has multiple oncogenic functions (Overholtzer et al., 2006). Moreover, increased expression and presence of YAP in the nucleus was found in different types of cancer, including colorectal cancer (Zhao et al., 2010). However, the underlying mechanisms regulating YAP in both normal and malignant epithelium remain largely unknown. It was shown that ECM composition and ECM stiffness plays an important role for the YAP activation (Gjorevski et al., 2016; Montagner and Dupont, 2020). Interestingly, YAP/TAZ deletion in a mouse model of colon tumorigenesis protects mice from tumor development (Azzolin et al., 2014), and there is a molecular evidence that YAP can be involved in tumor initiation in intestinal cancer (Taniguchi et al., 2017). Here, we show that α-SMA+ MFs, as one of the major producers of ECM, can regulate YAP in the intestinal epithelium, pointing to a crucial role of intestinal MFs as a constituent of the intestinal stem cell niche, and suggesting a potential role during tumor initiation. Future studies should aim at deciphering the detailed molecular mechanism including Yap upstream signals.

Finally, our work emphasizes the importance of the microenvironment in regulating epithelial plasticity. By different approaches such as various cellular assays, analysis of histology, assessment of cellular differentiation and gene expression profiling, we show that exposure to α-SMA+ MFs reversibly induces phenotypic convergence of wild type crypts to tumor crypts. Cellular plasticity in intestinal epithelial cells is thought to be dictated by the genetic interactions such a genetic inactivation of the Apc gene (Dow et al., 2015), and Apc restoration induces cellular differentiation and tumor regression. In contrast, our study unravels a potential mechanism of non-genetic control of epithelial plasticity, which could be mediated by α-SMA+ MFs and can be regulated by both intrinsic and extrinsic factors. As the number of α-SMA+ MFs is increased during intestinal tumorigenesis (Adegboyega et al., 2002) and α-SMA+ MFs can be used to predict disease relapse in colorectal cancer (Tsujino et al., 2007), future studies should consider intestinal α-SMA+ MFs as a potential biomarker for tumor initiation during colorectal cancer surveillance endoscopies.

One of the limitations of our study is that the inflammatory component, that is an important factor for tumor development in gastrointestinal tract, is missing in our 3D co-culture system. Since mouse infection with *Helicobacter pylori* results in upregulation of R-Spondin3, a potent Wnt signaling enhancer, in gastric myofibroblasts (Sigal et al., 2017), it would be interesting to recapitulate our findings using MF – epithelial - infectious agent - immune cell 3D co-culture model. Will canonical Wnt signaling be upregulated in epithelial cells in such a co-culture system?

To summarize, our study provides compelling evidence that the intestinal epithelium does not act autonomously but is regulated by MF-derived extrinsic factors. We show that intestinal α-SMA+ MFs can dictate the fate of intestinal epithelium and have impact on the expression of genes associated with cell cycle, cellular metabolism and tumor-initiation signaling pathways in the intestinal crypt, specifically through a TGF-β and YAP dependent signaling pathway.

## SUPPLEMENTARY FIGURE LEGENDS

**Figure S1.**
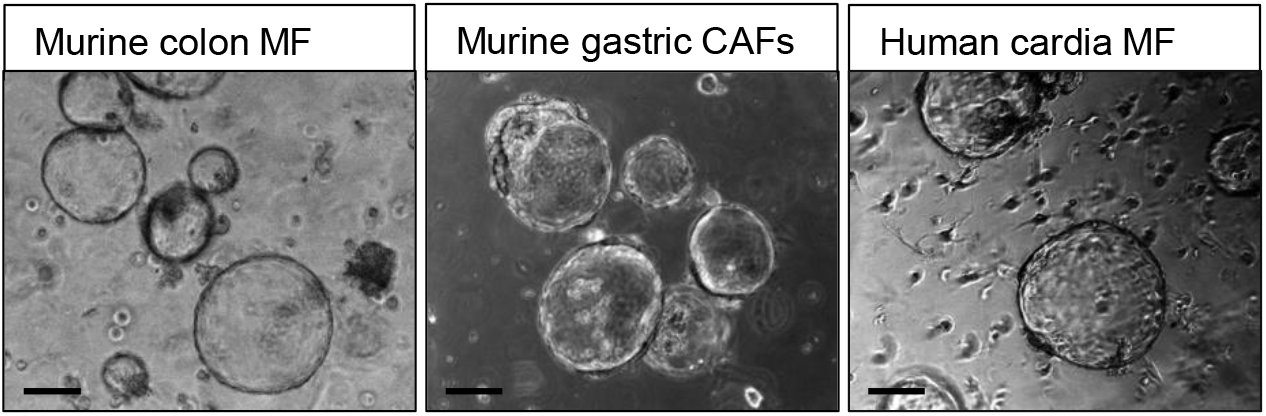
related to Figure 1.Morphology of enteroids in the presence of murine colon MFs, murine gastric carcinoma associated fibroblasts (CAFs) and human cardia MFs. Scale bar 100 μm.

**Figure S2.**
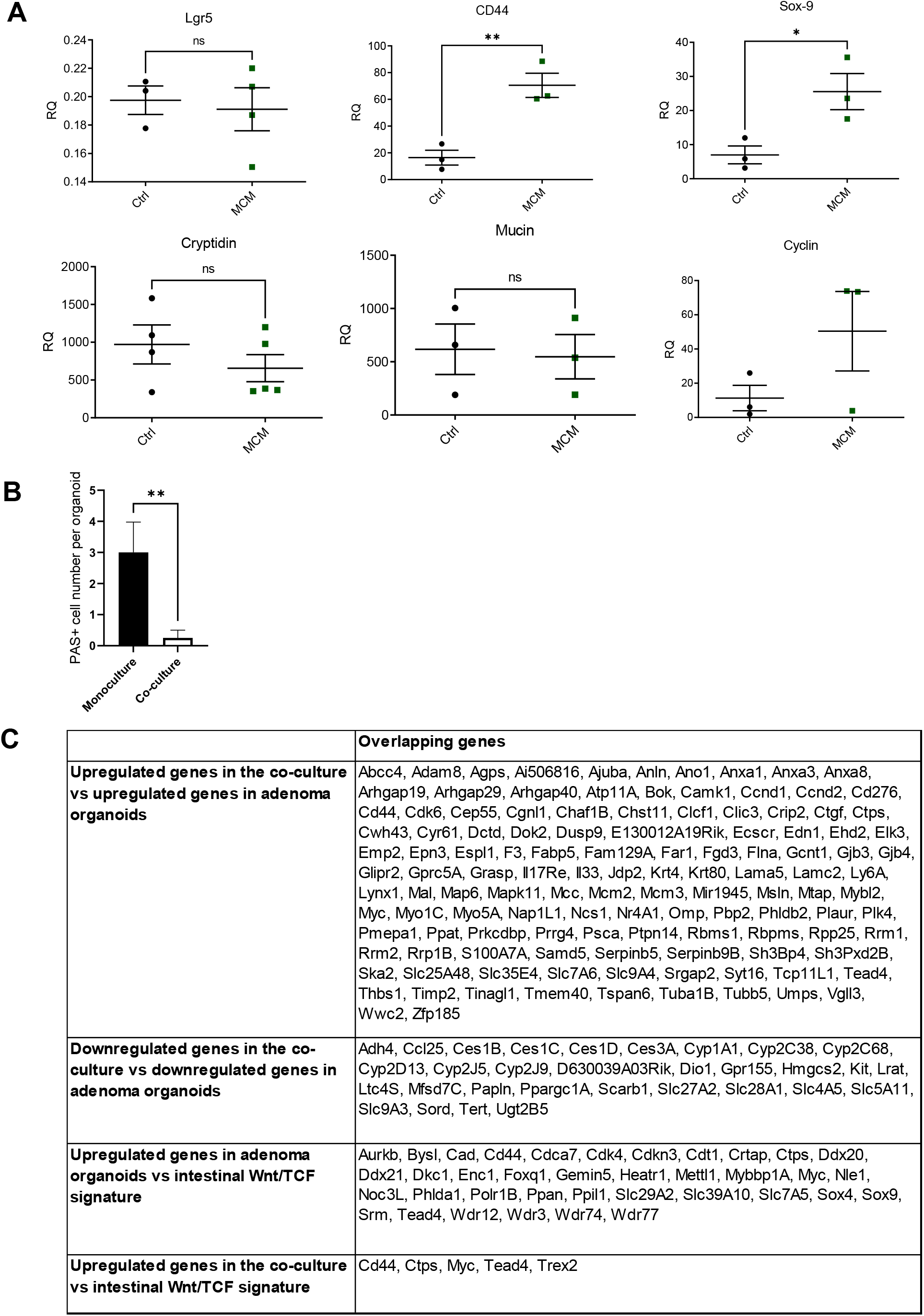
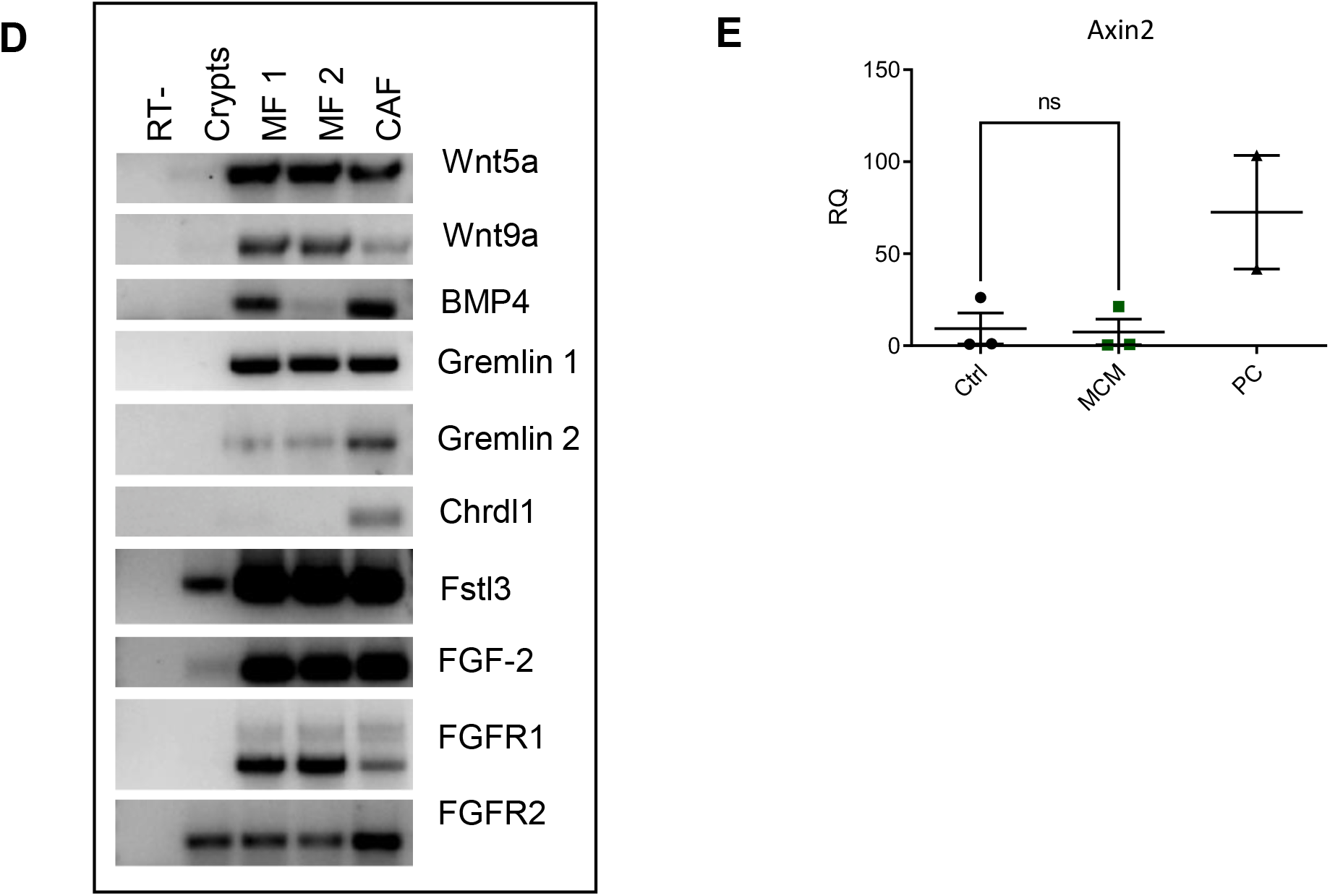
related to Figure 3.**A**. RTqPCR analysis of the expression of Lgr5, Sox-9 (p = 0.0348), CD44 (p = 0.0069), cyclin D1, mucin-2 and cryptidin-5 in the SI organoids incubated with myofibroblast conditioned medium (MCM). For the control (Ctrl), SI organoids were cultured in the same medium as used for the myofibroblast culture. Two-tailed t-test. **B**. Quantification of PAS+ cells in SI organoids cultured as monoculture and in the co-culture with SI myofibroblasts. Mean ± SEM, two-tailed t-test, p = 0.0032. **C**. Overlapping gene lists from microarray data. **D**. Expression of niche factors in the intestinal crypts cultured *in vitro* and stromal cells evaluated by RT-PCR. RT-, negative control for reverse transcription. MF, small intestinal myofibroblasts. CAF, carcinoma associated myofibroblasts. **E**. RTqPCR analysis of the expression of *Axin2* in the SI organoids treated with MCM. As positive control the tumor Apc^+/1638N^ enteroids were used.

**Figure S3.**
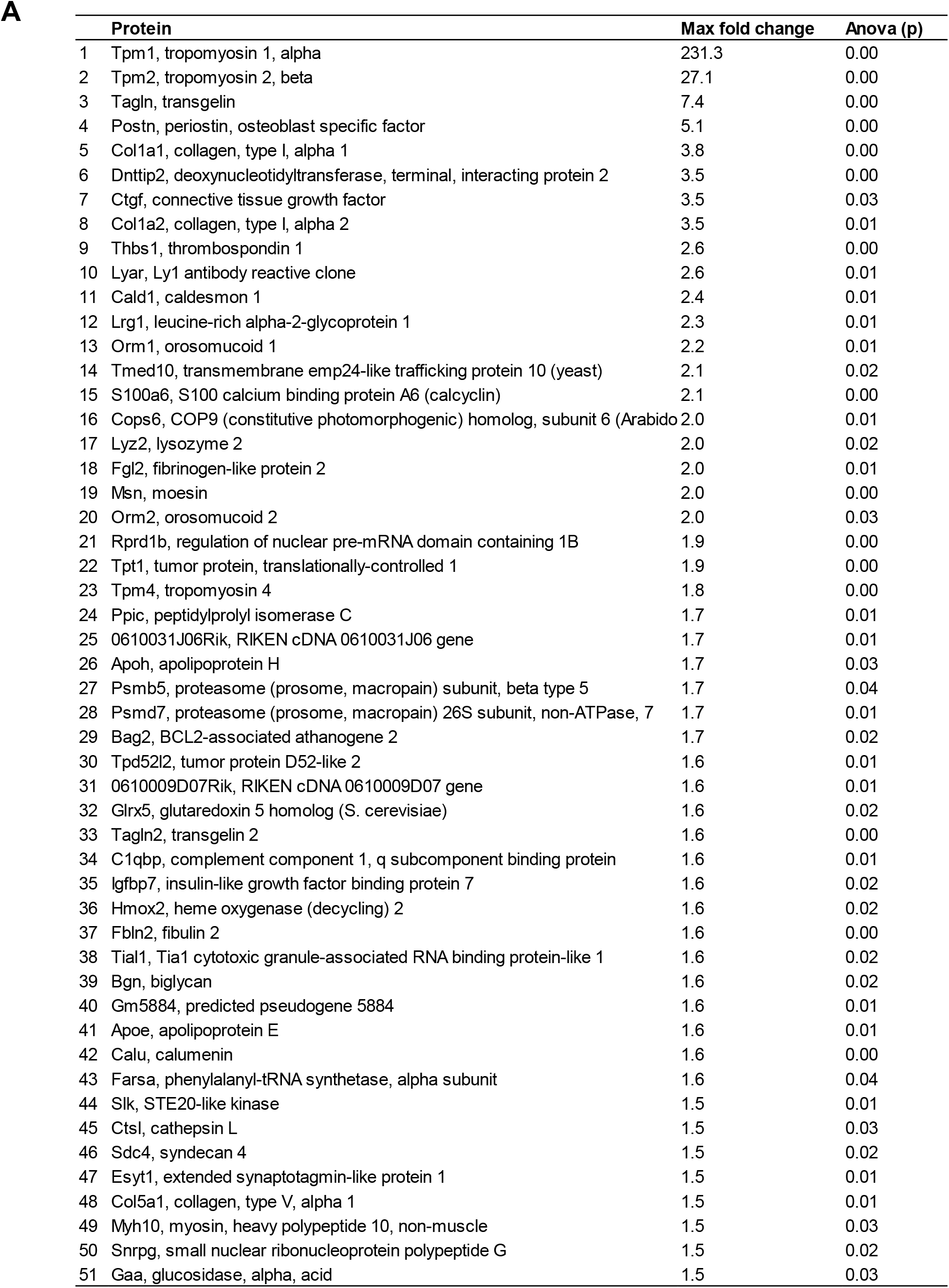
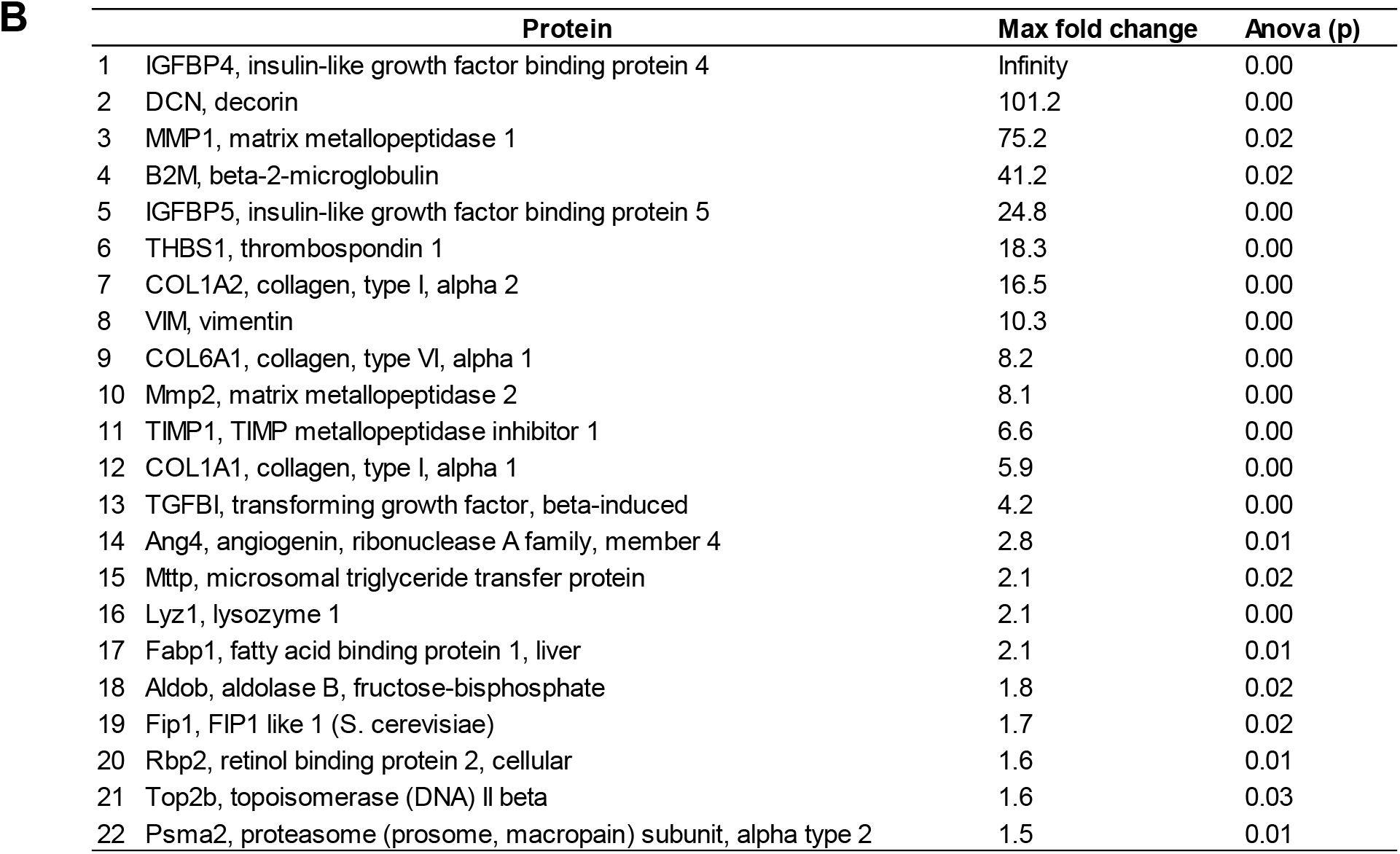
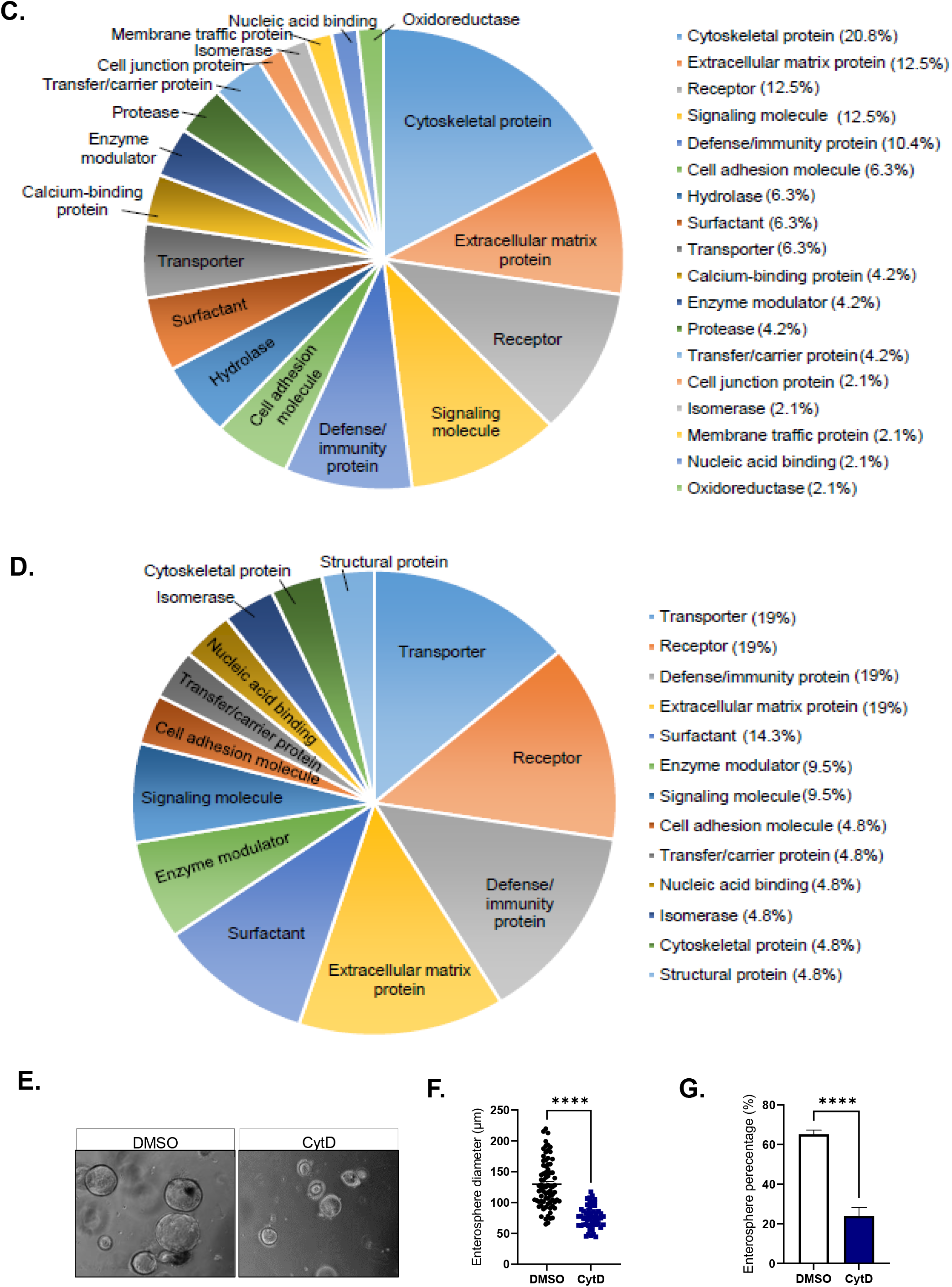
related to Figure 4.Mass spectrometry analysis of the supernatant from the stroma-epithelium co-cultures. **A**. Proteins significantly upregulated in the co-culture of murine small intestinal myofibroblasts and small intestinal crypts. **B**. Proteins significantly upregulated in the co-culture of human cardia myofibroblasts and small intestinal crypts. **C**. Protein class analysis of the significantly upregulated protein candidates in the co-culture of murine small intestinal myofibroblasts and small intestinal crypts. **D**. Protein class analysis of the significantly upregulated protein candidates in the co-culture of human cardia myofibroblasts and small intestinal crypts. **E**. Morphology of enteroids treated with myofibroblast conditioned medium (MCM) and cytochalasin D; scale bar 100 μm. **F**. Enterosphere diameter in enteroids treated with MCM and cytochalasin D. Two-tailed t-test, p < 0.0001. **G**. Enterosphere percentage in enteroids treated with MCM and cytochalasin D. Two-tailed t-test, p < 0.0001.

## EXPERIMENTAL PROCEDURES

### Isolation and culture of small intestinal myofibroblasts

Small intestinal (SI) myofibroblasts (MFs) were isolated from wild type (WT) C57BL/6 mice and cultured as previously described (Pastuła et al., 2016). Detailed procedure was described in the supplementary file. All animal studies and procedures were approved by the ethics committees at TUM.

### Isolation and culture of small intestinal crypts (small intestinal organoid culture, enteroids)

SI crypts were isolated from WT C57BL/6 mice and cultured in 3D culture system as previously described (Pastuła and Quante, 2016). Detailed procedure was described in the supplementary file.

### Direct co-culture studies

WT SI organoids and WT SI MFs were pelleted together and mixed with Matrigel, and 50 μl of the Matrigel/cell suspension was pipetted per well in a 24-well plate. Cells were cultured either in the presence or absence of R-Spondin, EGF and Noggin. After 24 h spheroid percentage was analyzed: (spheroid number/ total number of crypts) x 100%.

### Indirect co-culture studies

For an indirect co-culture SI MFs were seeded on the insert (Transwell, 24 mm diameter insert, 0.4 μm pore size, tissue culture treated polyester membrane, Corning) that was placed in a 6-well plate. Cells were cultured overnight in a medium composed of RPMI, 10% FBS and 1% penicillin/ streptomycin. Then, SI organoids were mixed with Matrigel and seeded in the 6-well plate, below the insert. Cells were cultured for 24 h in the Basal Medium. After that, organoids were harvested for RNA isolation.

### Human biopsies

For analysis of different stages of human colon carcinogenesis human tissue material from the Munich Biobank of the Department of Pathology was utilized to analyze normal colon mucosa (n=9), adenoma with low grade dysplasia (LGD) (n=9) or high-grade dysplasia (HGD) (n=4) and colon cancer (n=9).

### Ethics Approval

This study was approved by the TUM Ethics Committee (approval no. TUM 5291/12 and TUM 5428/12).

### RNA isolation, RT-PCR and real-time PCR

RNA was isolated using RNeasy mini Qiagen kit according to the manufacturer’s guidelines. The reverse transcription - PCR (RT-PCR) was conducted using GoTaq® Green Master Mix, 2X (Promega) according to manufacturer’s instructions. The RT-PCR products were analyzed on a 1.5% agarose gel. For the real-time PCR 2x QuantiFast SYBR Green PCR Master Mix (Qiagen) was used. The reaction was performed with LightCycler 480 Roche according to manufacturer’s instructions. As a reference gene GAPDH was used. Primer sequences for RT-PCR and real-time PCR are listed in the supplementary file.

### Microarray analysis

For the microarray analysis the in-direct co-culture system was applied (described above). In addition to the WT SI organoid monoculture, WT SI organoid – SI MF co-culture, microarray for the adenoma organoids was performed. After 24 h epithelial cells were harvested for RNA analysis. RNA was isolated with RNeasy mini Qiagen kit and afterwards ethanol precipitated. Hybridization to the Affymetrix Mouse Gene 2.1 ST Array Plates was done at the Kompetenzzentrum Fluoreszente Bioanalytik der Universität Regensburg. Data was normalized with RMA (Irizarry et al., 2003), as implemented in Oligo (Carvalho and Irizarry, 2010). Data was deposited in the Gene Expression Omnibus (Barrett et al., 2012) (Accession GSE124684). Differential expression was analyzed with weighted Limma (Ritchie et al., 2015), using a significance cutoff of the Benjamini-Hochberg false discovery rate, fdr ≤ 0.05 (Benjamini and Hochberg, 1995). Microarray data were analyzed with the DAVID search tool (Huang et al., 2008) applied to the Bioinformatics/ KEGG pathway database (Kanehisa et al., 2004) and the PANTHER search tool and database (Huang et al., 2008).

## AUTHOR CONTRIBUTIONS

A.P. designed and conducted experiments, analyzed data, and wrote the manuscript. K-P. J., U.E, M. R-F., M. B. and M. F. performed experiments. R. M. S. and T. C. W. analyzed data. S. H. performed mass spectrometric analysis. R. A. F. analyzed microarray data. J. S-H. and K. S. performed IHC and provided human colon samples. MQ supervised the study, evaluated the manuscript and financed the studies.

## ACKNOWLEDGEMENTS

A.P. received Laura Bassi-Award (TUM) and was supported by the Dr.-Ing. Leonhard-Lorenz-Stiftung (TUM). AP is currently supported by the National Science Centre in Poland (NCN) grant (2020/39/D/NZ5/03552). We thank the tissue bank of TUM and Klinikum rechts der Isar and the Comparative Experimental Pathology (School of Medicine, TUM) for excellent technical support. M.Q. received funding from the Max Eder Program of the Deutsche Krebshilfe.

## Graphical summary

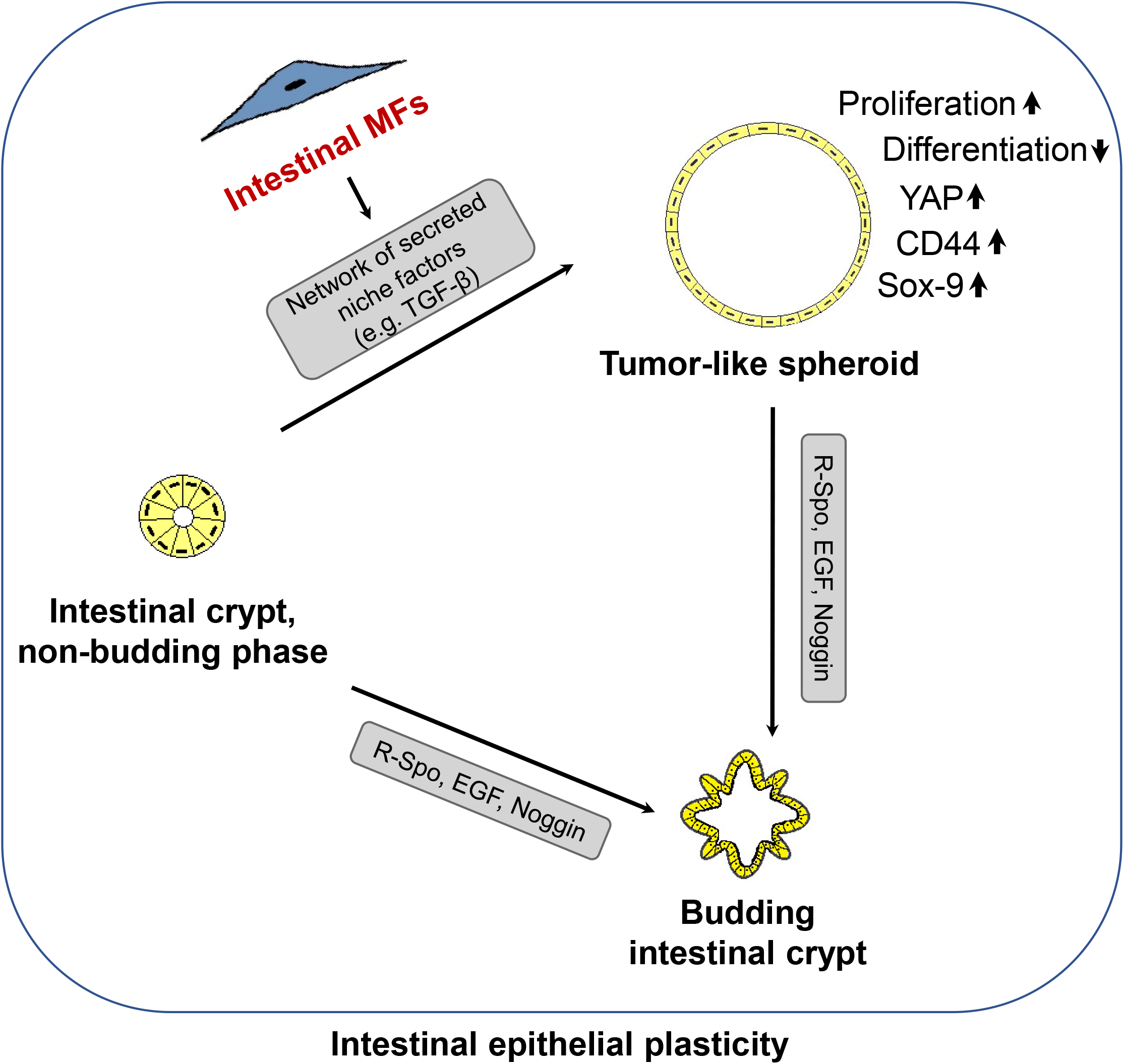

